# A chemotactic sensor controls *Salmonella*-host cell interaction

**DOI:** 10.1101/2021.05.14.443827

**Authors:** Stefanie Hoffmann, Kathrin Gendera, Christiane Schmidt, Peter Kirchweger, Axel Imhof, Christian Bogdan, Yves A. Muller, Michael Hensel, Roman G. Gerlach

**Affiliations:** Project Group 5, Robert Koch Institute, Wernigerode, Germany; Division of Biotechnology, Department of Biology, Friedrich-Alexander-Universität Erlangen-Nürnberg (FAU), Erlangen, Germany; BioMedical Center and Center for Integrated Protein Sciences Munich, Ludwig-Maximilians-Universität München, München, Germany; Mikrobiologisches Institut – Klinische Mikrobiologie, Immunologie und Hygiene, Universitätsklinikum Erlangen and Friedrich-Alexander-Universität Erlangen-Nürnberg, Erlangen, Germany; Abteilung Mikrobiologie, Universität Osnabrück, Osnabrück, Germany and CellNanOs - Center for Cellular Nanoanalytics Osnabrück, Universität Osnabrück, Osnabrück, Germany

**Author notes:** Corresponding author: Roman G. Gerlach **Email:**. Department of Chemical and Structural Biology, Faculty of Chemistry, Weizmann Institute of Science, Rehovot, Israel (Peter Kirchweger). **Author Contributions:** R.G.G., S.H., M.H. and Y.A.M. designed research; S.H., K.G., C.S. and P.K. performed research; R.G.G., S.H. and A.I. analyzed the data; and R.G.G., C.B., M.H. and Y.A.M. wrote the paper.

**Keywords:** Salmonella, chemotaxis, adhesion, aspartate

## Abstract

Intimate cell contact and subsequent type three secretion system-dependent cell invasion are key steps in host colonization of *Salmonella*. Adhesion to complex glycostructures at the apical membrane of polarized cells is mediated by the giant adhesin SiiE. This protein is secreted by a type 1 secretion system (T1SS) and needs to be retained at the bacterial surface to exert its adhesive function. Here, we show that SiiE surface expression was linked to the presence of L-aspartate sensed by the *Salmonella*-specific methyl-accepting chemotaxis protein CheM. Bacteria lacking CheM were attenuated for invasion of polarized cells, whereas increased invasion was seen with *Salmonella* exposed to the non-metabolizable aspartate analog α-methyl-D, L-aspartate (MeAsp). While components of the chemotaxis phosphorelay or functional flagella were dispensable for the increased invasion, CheM directly interacted with proteins associated with the SiiE T1SS arguing for a novel non-canonical signaling mechanism. As a result, CheM attractant signaling caused a shift from secreted to surface-retained and adhesion-competent SiiE. Thus, CheM controls the virulence function of SiiE in a precise spatio-temporal fashion depending on the host micro-milieu.

## INTRODUCTION

Many pathogenic bacteria strongly rely on their ability to get into close contact to eukaryotic host cell surfaces. By means of different adhesins, they are able to colonize mucosal surfaces or invade cells and establish their niches in host organisms. *Salmonella enterica* subsp. *enterica* serovar Typhimurium (STM) is a pathogen that is capable to infect diverse hosts and usually causes a self-limiting gastrointestinal infection. STM can invade non-phagocytic cells by deploying a type III secretion system (T3SS) that is encoded by the *Salmonella* pathogenicity island 1 (SPI-1) (1). An intimate contact between the pathogen and the host cell is essential for the subsequent translocation of effector molecules by the SPI-1-dependent T3SS (T3SS-1). This triggers an inflammatory host immune response, which does not only weaken the enterocyte barrier function, but also helps STM to compete with the intestinal microbiota (2). While the T3SS-1 itself can already mediate adhesion (3), additional adhesive structures such as Fim fimbriae (4) or the giant non-fimbrial adhesin SiiE (5) are also critical for bacterial attachment, depending on the type of host cell. Transcriptional co-regulation of SPI-1 with the SiiE-encoding *Salmonella* pathogenicity island 4 (SPI-4) is the basis of this functional cooperation (5, 6).

In line with previous findings on other polarized cells (7), it was recently shown that apical invasion of intestinal epithelial cells requires SiiE (8). SiiE likely functions as a lectin recognizing glycostructures with terminal *N*-acetylglucosamine (GlcNAc) and/or α 2,3-linked sialic acid residues (9). The unique structural features of the ∼175 nm long 600 kDa adhesin SiiE allow *Salmonella* to overcome the anti-adhesive barrier function of the transmembrane, epithelial mucin MUC1, the extracellular domain of which is heavily decorated with O-linked glycans terminating in negatively charged sialic acids (8). SiiE comprises 53 bacterial immunoglobulin-like (BIg) domains that show distinct Ca^2+^ binding motifs crucial for its rigid tertiary structure (10, 11). SiiE is the only known substrate of the SPI-4-endoced T1SS and contains a complex C-terminal secretion signal (7). Based on similarity to other T1SS, e.g., the *E. coli* hemolysin system (12), secretion is likely achieved in a single step without a periplasmic intermediate. Considering these structural features, SiiE is thought to be permanently secreted into the extracellular space. A recent study showed that secreted SiiE suppressed the humoral immune response against *Salmonella* by reducing the number of IgG-secreting plasma cells in the bone marrow. Mechanistically, an N-terminal region of SiiE with high similarity to laminin β1 bound to β1 integrin (CD29) on IgG^+^ plasma cells, thereby preventing their interaction with laminin β1^+^CXCL12^+^ stromal cells which otherwise form a survival niche for plasma cells in the bone marrow (13). However, in line with its function as an adhesin, the protein was also found temporally retained on the surface of *Salmonella* (7, 8, 14). Nonetheless, it is still unclear how the surface expression of SiiE and hence the switch between its function as adhesin vs. immunosuppressant is regulated.

Here, we show that the presence of aspartate (Asp) promotes the surface expression of SiiE and the adhesion of *Salmonella*, which turned out to be dependent on the *Salmonella*-specific methyl-accepting chemotaxis protein (MCP) CheM, an ortholog of *E. coli* Tar/MCP-II. Using mass spectrometry, CheM and other MCPs were identified as interaction partners of the SPI-4 encoded SiiA and SiiB. SiiAB are associated with the T1SS and form an inner membrane proton (H^+^) channel with similarities to the ExbB/TolQ and MotAB family (15, 16). Using a set of consecutive MCP deletion mutants, we found that invasion of polarized cells by STM was attenuated upon deletion of *cheM*. Binding of Asp to CheM usually triggers bacterial chemotaxis towards the attractant gradient (17). We discovered that the addition of a non-metabolizable aspartate analog, α-methyl-D,L-aspartate (MeAsp), elevated host cell invasion by STM. Using mutants lacking downstream components of the chemotaxis phosphorelay or functional flagella, we observed that neither classical chemotaxis signaling nor bacterial motility contributed to the increased invasion. Instead, attractant-stimulation of CheM caused a shift from secreted to surface-retained and adhesion-competent SiiE. We therefore suggest that aspartate acts as microenvironmental cue that elicits the SPI-4-dependent adhesion to and invasion of polarized epithelial cells by *Salmonella* through a novel, non-canonical signaling pathway of the MCP CheM via SiiAB to the SPI-4 T1SS and SiiE.

## RESULTS

### SiiAB interact with CheM

Our previous results suggested that SiiE-mediated adhesion depends on the function of the SPI-4 T1SS-associated SiiAB proton channel (15). We set out to identify protein interaction partners as potential regulators of SiiA and/ or SiiB. Epitope-tagged SiiA or SiiB was expressed from low copy-number plasmids and used as bait proteins to purify complexes after crosslinking. Composition of complexes was determined by liquid chromatography coupled to mass spectrometry (LC-MS/MS). As expected, both bait proteins were under the top five identified proteins (Fig 1A). Moreover, our data confirmed the previously found (15) interaction between SiiA and SiiB, since both proteins were identified as prey while using the other as bait. Interestingly, a set of MCPs (Aer, Trg, McpB, McpC and CheM) was enriched in the SiiA complex while in the SiiB complex the MCP CheM was identified. MCPs act as sensors controlling flagellar movement towards attractants and away from repellants (18). In the following, we focused on CheM because the protein was previously implicated to have a role in *Salmonella* invasion of HeLa cells (19, 20) and appears to interact with both SiiA and SiiB. CheM is a *Salmonella*-specific MCP and an ortholog of *E. coli* Tar, but shares only 79% of sequence identity. Like known for Tar, the amino acid L-aspartate functions as an attractant stimulus for CheM while Co^2+^ and Ni^2+^ ions act as repellants (17, 21). However, in contrast to Tar, CheM does not respond to maltose (22).

**Figure 1.**
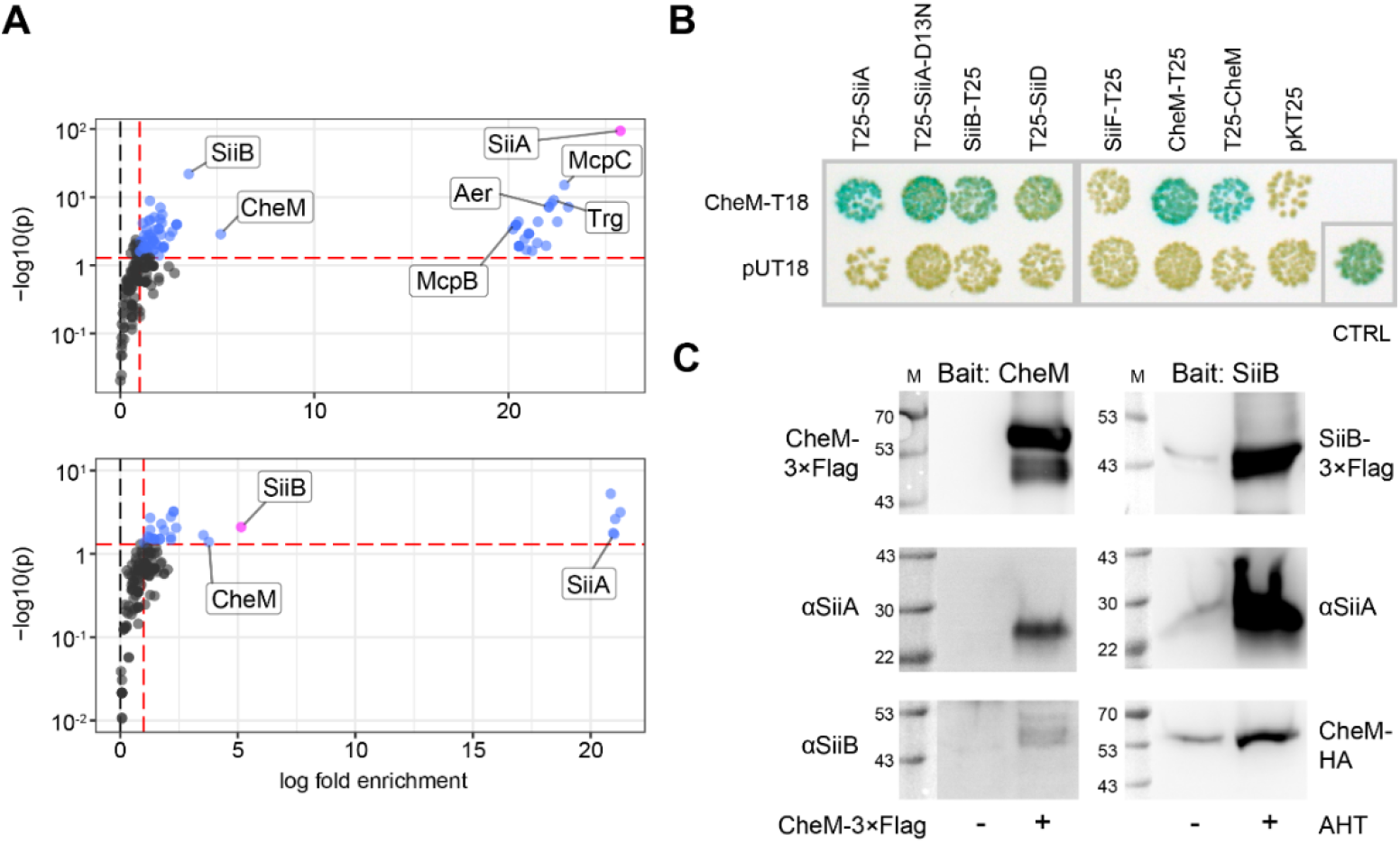
SPI-4 components interact with CheM. (A) Analysis of affinity purification mass spectrometry data using SiiA-3×Flag (upper panel) or SiiB-3×Flag (lower panel) as bait proteins (magenta dots). Red dashed lines show limits of significant enrichment (>2-fold, *p* < 0.05). Interacting proteins within these limits are depicted in blue with SPI-4 components and MCPs labeled. Summarized data of three independent experiments are shown. (B) Bacterial two hybrid assays evaluating the interaction between the T18 fragment of CyaA alone (pUT18, negative control) or fused to the C-terminus of CheM (CheM-T18) with the CyaA T25 fragment alone (pKT25, negative control) or T25 fused to the indicated SPI-4 proteins or CheM. Functional reconstitution of CyaA activity through protein-protein interactions resulted in blue color of the *E. coli* BTH101 reporter strain colonies. A positive control (CTRL) was included based on the interaction of GCN4 leucine zippers. (C) Co-immunoprecipitation using CheM-3×Flag (left panels) or SiiB-3×Flag (right panels) as bait proteins. A plasmid-encoded copy of *cheM* was expressed from its natural promoter either without (left lane) or with (right lane) 3×Flag epitope tag. SiiA and SiiB were detected using polyclonal antibodies. Expression of SiiAB-3×Flag from plasmid pWRG905 was induced with addition of 50 ng/mL anhydrotetracycline (AHT) or left uninduced (left lane). While SiiA was detected with a specific antiserum, a plasmid encoding for CheM-HA was co-transformed allowing CheM detection via HA tag. M = molecular weight marker with protein sizes in kDa.

To confirm the interactions between SiiAB and CheM identified by MS, we performed a bacterial two-hybrid (B2H) assay which is based on the functional complementation of *Bordetella pertussis* adenylate cyclase (CyaA) from T25 and T18 fragments (23). The T25 fragment was fused to SiiA, SiiB, SiiD, SiiF or to a non-functional SiiA^D13N^ mutant (15), while the T18 fragment was fused to CheM. Blue colonies of the *cyaA*-deficient *E. coli* reporter strain BTH101 (24) indicated functional CyaA protein complementation and thus protein-protein interaction. We observed a strong interaction (dark blue colonies) between CheM-T18 and T25 fusions of SiiA and SiiA-D13N (Fig 1B). Moreover, high reporter activity was also observed when testing for the known dimerization of Tar/CheM (18) with co-expression of CheM-T18 and T25-CheM or CheM-T25 (Fig 1B). Lighter blue colonies were observed for the co-expression of CheM-T18 and SiiB-T25 or T25-SiiD showing β-galactosidase activity comparable to that of the positive control (Fig 1B). Furthermore, we performed co-immunopecipitation (co-IP) using epitope tagged proteins. When CheM-3×Flag was used as bait, both SiiA and SiiB were identified as prey proteins. Vice versa, using SiiB-3×Flag as bait, SiiA and epitope-tagged CheM-HA were detected as interaction partners (Fig 1C). Thus, using three independent approaches, we established that CheM interacts with both SiiA and SiiB while confirming the known SiiAB complex (15) and CheM dimerization (18).

### Role of MCPs for invasion of polarized MDCK

Bacterial motility is required for efficient invasion of *Salmonella* into HeLa (25) and polarized Caco-2 cells (26). In a previously published study, we further assessed the impact of chemotaxis on invasion efficiency of non-polarized HeLa cells using sequential deletion of up to seven MCP genes (19). We found that loss of CheY or CheM led to an increase of *Salmonella* invasion (19) as observed earlier for smooth swimming mutants (20, 27). Because SPI-4 function was shown to play a role for adhesion to polarized cells only (5, 7, 8), we aimed to investigate the role of individual MCPs on *Salmonella* invasion of polarized Madin-Carby Canine Kidney (MDCK) cells. Host cells were infected with STM wild-type (WT), a smooth swimming *cheY* mutant, Δ*siiF* (non-functional SPI-4 T1SS) and MCP mutant strains as described (19), followed by quantification of intracellular bacteria and subsequent normalization to STM WT. Interestingly, all MCP mutants missing the *cheM* gene exhibited reduced invasion in polarized MDCK cells. In contrast, elevated invasion rates were observed for the same *cheM*-lacking mutants when using non-polarized HeLa cells as described before (19) (Fig 2). While a smooth swimming Δ*cheY* strain showed a 2-fold increased invasion rate in HeLa, the mutant was significantly attenuated in MDCK arguing for a role of chemotaxis for efficient invasion of polarized cells. In line with the known importance of SPI-4 for the adhesion to and invasion of MDCK cells (7), very few intracellular bacteria harboring an E627Q mutation within the Walker B motif of the SiiF ABC protein were detected (Fig 2). Thus, the type of infection model (non-polarized vs. polarized cells) determine the impact of CheM function on STM invasion which mirrors the differences seen for SPI-4 function (7).

**Fig 2.**
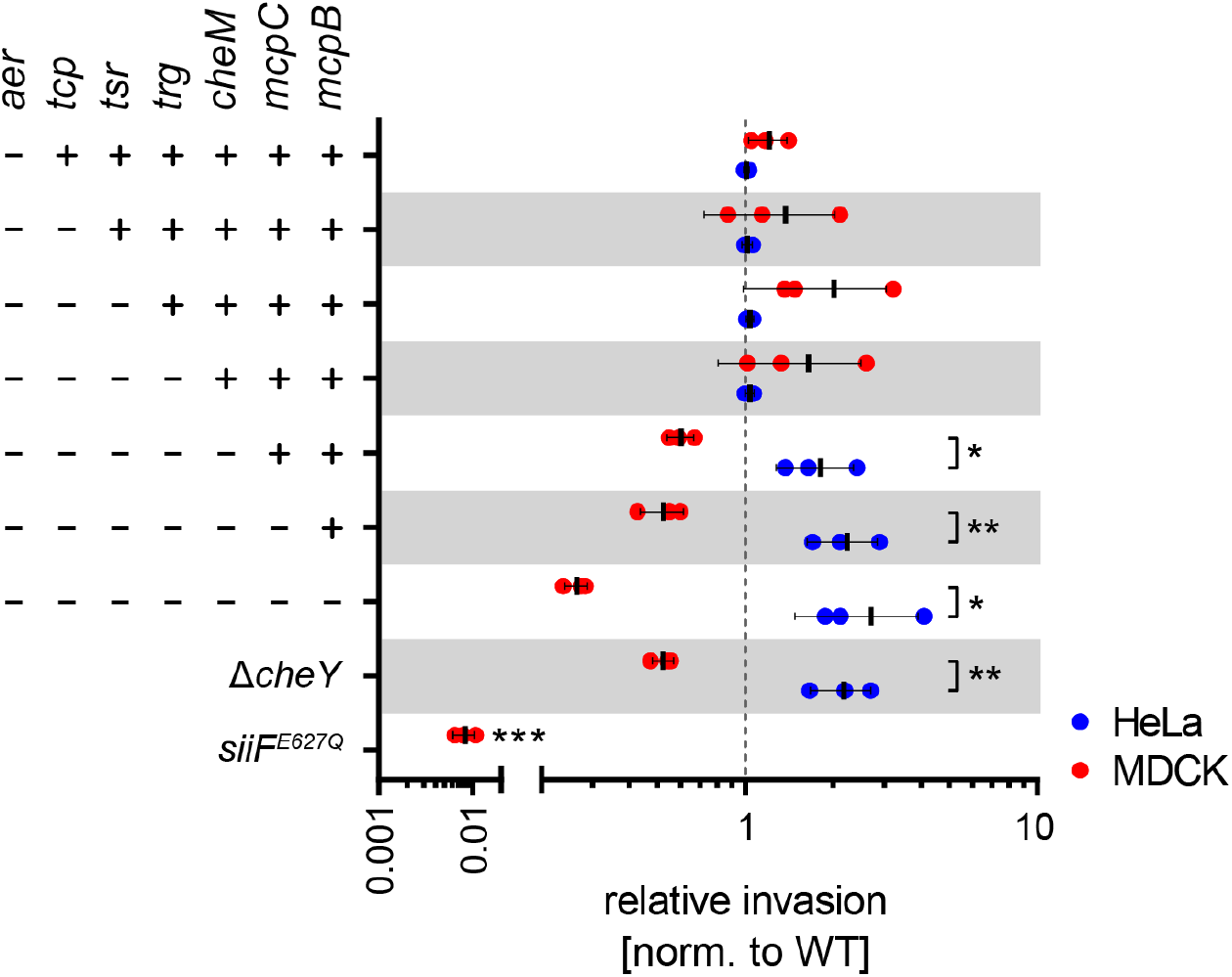
Role of MCPs for bacterial invasion. Relative invasion rates as normalized to *S*. Typhimurium (STM) wild-type (WT, dotted line) after one hour of infection of indicated sequential MCP deletion strains into HeLa (blue) and MDCK (red) cells are shown. The non-chemotactic Δ*cheY* strain and a *siiF* E627Q mutant with a non-functional SPI-4 were included as controls. Data of three independent experiments done in triplicates are depicted. Statistical significance was calculated using unpaired, two-tailed *t* test between groups or a one sample *t* test against the hypothetical value 1 (*siiF* E627Q) and were defined as * for *p* < 0.05 and ** for *p* < 0.01 and *** for *p* < 0.001.

### CheM attractant binding fosters *Salmonella* invasion of polarized cells

To characterize a possible functional link of CheM to the SPI-4-encoded T1SS, we constructed two low-copy-number plasmids that encode *Salmonella cheM* or, as a control, *E. coli tar*, each modified with a C-terminal 3×Flag-tag under control of the STM *cheM* promoter (P_*cheM*_). After introducing the plasmids in a mutant lacking all seven MCP genes (Δ7 MCP) (19), Western blot demonstrated similar expression of both proteins with the cytosolic protein DnaK as loading control (Fig 3A). Next, Δ7 MCP was transformed with the empty vector control (pWSK29) and plasmids encoding for CheM (pCheM) or Tar (pTar) without epitope tag and these strains were further functionally characterized in swarming assays using soft agar plates. While the strain harboring pWSK29 did not swarm, pCheM and pTar conferred swarming ability to the mutant. However, compared to STM WT (> 5 cm, not shown), both plasmid-complemented Δ7 MCP showed a reduced swarming distance with ∼4 cm (pCheM) and ∼1 cm (pTar), respectively (Fig 3B). To test more specifically the ability to respond to CheM attractants, we performed a capillary assay as described by Adler (28) using MeAsp (29) (Fig 3C, left panel). Quantifying the bacteria within the fixed-volume capillary revealed significantly more cells in case of the pCheM-complemented strain, compared not only to the vector control, but also compared to WT (Fig 3C, right panel).

**Fig 3.**
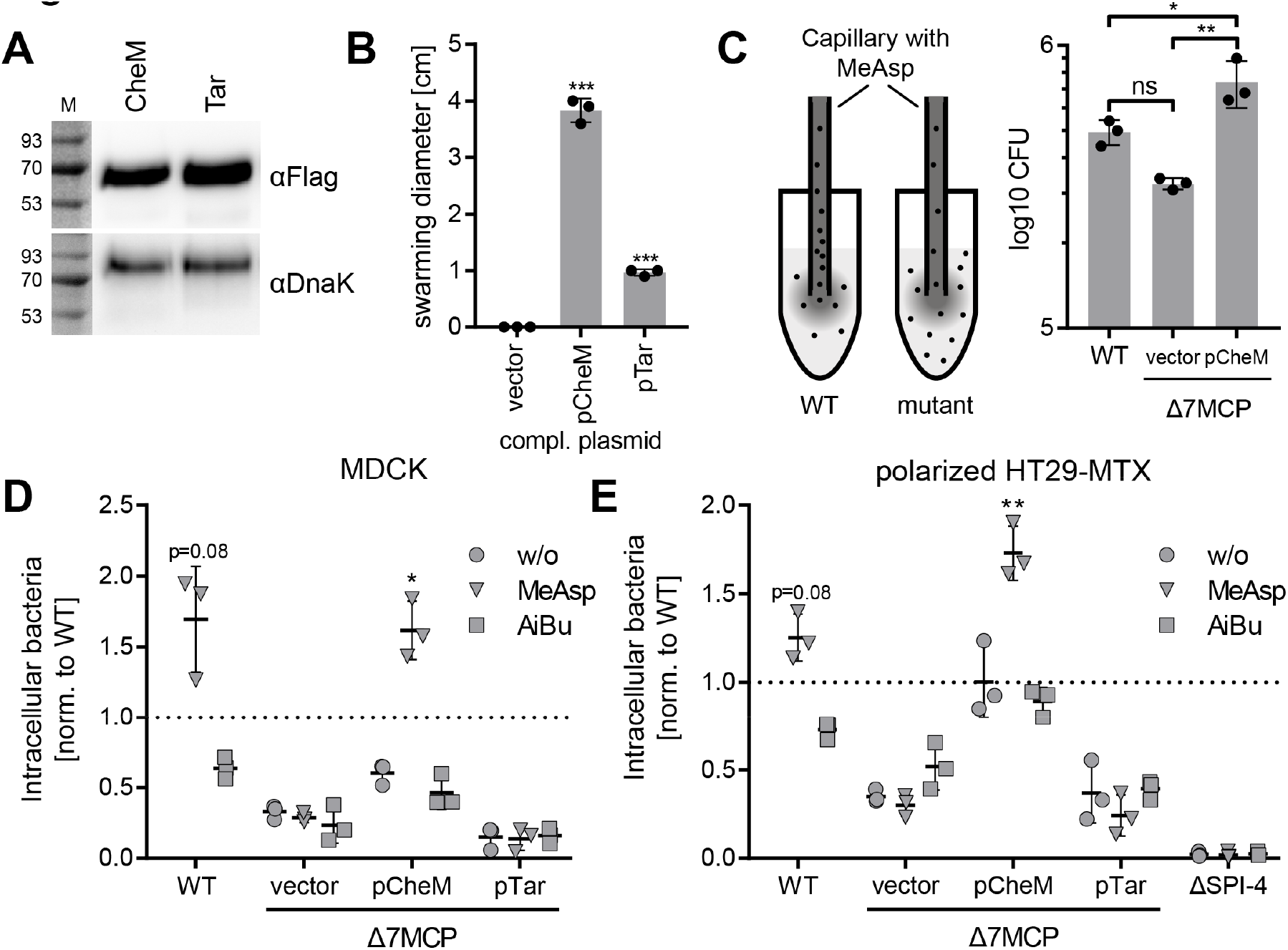
CheM complementation and impact of CheM signaling on invasion of polarized cells. (A) Expression of either CheM-3×Flag or Tar-3×Flag from the CheM promoter in a *S*. Typhimurium (STM) strain lacking all 7 MCP genes (Δ7 MCP) was detected with a Flag-specific monoclonal antibody. Equal sample loading was demonstrated by DnaK. M = molecular weight marker with protein sizes in kDa. (B) The Δ7 MCP strain transformed with the empty vector pWSK29 (vector) or vectors encoding for CheM (pCheM) or Tar (pTar) were subjected to swarming assays on soft agar plates. Mean swarming diameters ± SD after 8.5 h of growth are depicted for three independent experiments. Statistical significance was calculated using a one sample *t* test against the hypothetical value 0 (vector control) and were defined as *** for *p* < 0.001. (C) Principle of capillary assay. Wild-type (WT) bacteria with functional CheM-dependent chemotaxis swim towards a gradient of α-methyl-D, L-aspartate (MeAsp) generated by an attractant-filled capillary. No enrichment within the capillary is observed for mutants with defects in CheM-signaling (left panel). Mean amount ± SD of STM WT or the Δ7 MCP with empty vector (vector) or pCheM within the capillary after 1 h. of chemotactic movement from three independent experiments are shown (right panel). One way ANOVA with Tukey’s multiple comparison test was calculated and was defined as * for adj. *p* < 0.05 and ** for adj. *p* < 0.01. (D and E) Relative invasion rates as normalized to STM WT (black dotted line) after one hour of infection of the indicated strains into polarized MDCK (D) or HT29-MTX (E) cells are shown. The strains were either grown without (w/o) attractant or with addition of 10 mM MeAsp or 10 mM AiBu. A SPI-4 deficient strain (ΔSPI-4) was included as control for HT29-MTX. Mean ± SD from three independent experiments are depicted. Statistical significance of strains with increased invasiveness was calculated using a one sample *t* test against the hypothetical value 1 and were defined as * for *p* < 0.05 and ** for *p* < 0.01.

We hypothesized that not the CheM protein itself, but rather CheM signaling elicited by the binding of CheM ligands (i.e. attractants) may have an impact on SPI-4 function and subsequently on invasion of polarized epithelial cells. Usually, attractant binding inactivates autokinase activity of MCP-coupled CheA, thus reducing phosphoryl transfer to the response regulators CheY and CheB. While low CheY∼P results in counter-clockwise (CCW) flagellar rotation and straight swimming, receptor methylation is high due to low CheB∼P methylesterase activity (18). Therefore, STM WT and the Δ7 MCP mutant containing either the vector control, pCheM or pTar were tested for invasion of MDCK without attractant, in the presence of 10 mM MeAsp or, as a control, 10 mM of the non-metabolizable Tsr attractant α-aminoisobutyrate (AiBu) (30, 31). While AiBu had no or, in case of STM WT, even a detrimental effect on invasion, addition of MeAsp elevated invasion capability of WT and pCheM-complemented Δ7 MCP (Fig 3D). The pCheM vector partially complemented the invasion defect of the Δ7 MCP mutant in the absence of attractant or with addition of AiBu, while the strains carrying pTar or the vector control were attenuated for invasion regardless of attractant supplementation (Fig 3D).

To verify our findings obtained with MDCK cells, we employed HT29-MTX cells (8, 32) as an alternative infection model. In contrast to non-polarized 1-day cultures (Fig S1A, upper panel), polarized monolayers with significant amounts of mucus were formed after 21 days of culture (Fig S1A, lower panel). Similar to HeLa cells (7), *Salmonella* invasion of non-polarized HT29-MTX cells required T3SS-1 but was independent of SPI-4 (Fig S1B). Invasion of polarized HT29-MTX cells, however, was strongly dependent on an intact SPI-4 locus (Fig 3E) as observed before (8). In close accordance with the MDCK data, elevated invasion of HT29-MTX cells was observed for WT and Δ7 MCP [pCheM] in the presence of MeAsp (Fig 3E). However, pCheM was able to complement the Δ7 MCP mutant to the level of WT STM without attractant or with 10 mM AiBu. In contrast, low invasion rates were observed for cells without CheM (Fig 3E).

Taken together, using two infection models based on polarized cells, we observed a stimulating effect of the CheM ligand MeAsp on *Salmonella* invasion. The phenotype was specifically dependent on the presence of CheM. No increase in invasion was seen in strains only expressing *E. coli* Tar or with addition of the Tsr ligand AiBu.

### Augmented invasion after CheM stimulation is independent of motility and chemotaxis

Bacterial motility and chemotaxis towards energy sources was shown to be required for *Salmonella* virulence *in vivo* (33-35). Because our data also suggest a promoting effect of chemotaxis for invasion of polarized cells, we set out to characterize the role of motility and the chemotaxis phosphorelay pathway for the observed phenotype in more detail. We generated a non-motile mutant lacking the flagella-specific ATPase *fliI* and employed, besides the *cheY*-deficient strain, a mutant lacking the histidine autokinase CheA. Together with MCPs and the coupling protein CheW, CheA dimers form a ternary complex that is the minimum requirement for chemosensing (18, 36). The *cheA* and *cheY* mutations were further combined with the Δ7 MCP mutant. These mutant strains were all attenuated for invasion of MDCK. Interestingly, the non-motile Δ*fliI* and the two 8-fold mutants (Δ7 MCP plus Δ*cheA* and Δ7 MCP plus Δ*cheY*) exhibited an even stronger phenotype with almost no invasion detectable (Fig 4A). While significantly more intracellular WT STM bacteria were found when grown in the presence of MeAsp, the mutants responded neither to this attractant nor to AiBu (Fig 4A).

**Fig 4.**
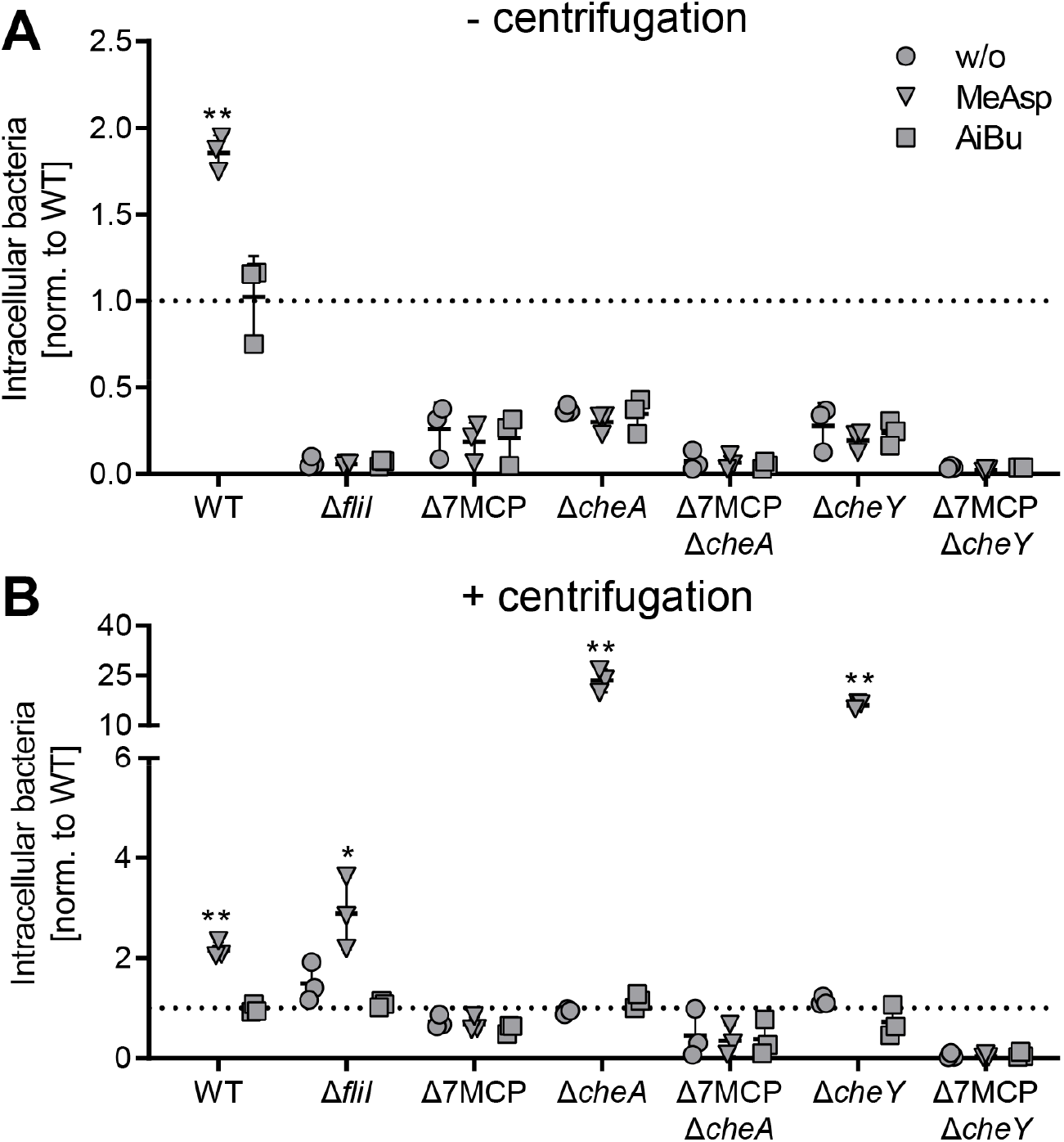
Impact of motility and chemotaxis components on *Salmonella* invasion of polarized MDCK. *S*. Typhimurium (STM) wild-type (WT), the non-motile Δ*fliI* mutant, the Δ7 MCP deletion strain, the chemotaxis mutants Δ*cheA* and Δ*cheY* or strains lacking besides *cheA* or *cheY* additionally all 7 MCPs were grown without attractant (w/o), in the presence of 10 mM MeAsp or 10 mM AiBu. Inoculi were added to the MDCK cells (A) or bacteria were brought in close host cell contact through centrifugation to compensate for lack of chemotaxis or motility (B). Intracellular bacteria were quantified after one hour of infection and relative invasion rates were calculated based on STM WT without attractant. Data of three independent experiments done in triplicates are depicted. Statistical significance of strains with increased invasiveness was calculated using a one sample *t* test against the hypothetical value 1.0 and was defined as * for *p* < 0.05 and ** for *p* < 0.01.

Motile bacteria exhibit an invasion advantage due to near surface swimming and thus higher probability to encounter a host cell (25). To compensate for the lack of motility, we used centrifugation to bring bacteria into close proximity to the host cells, which allows investigating bacterial invasion following adhesion despite the absence of bacterial motility. Under these conditions, the Δ*fliI* mutant behaved like WT with significantly increased invasion in the presence of MeAsp (Fig 4B). The invasion capability of the Δ7 MCP mutant was not altered by centrifugation. While Δ*cheA* and Δ*cheY* mutants behaved similar to WT bacteria without attractant or with AiBu, they showed vastly increased invasion rates when MeAsp was added (Fig 4B). In contrast, *Salmonella* with a *cheA* or *cheY* mutation and simultaneous deletion of all MCPs (ΔcheA Δ7 MCP or Δ*cheY* Δ7 MCP) lost the responsiveness to MeAsp and the ability to invade host cells. Thus, MeAsp fosters *Salmonella* invasion in a CheM-dependent manner, but this process is independent of bacterial motility and the chemotaxis phosphorelay pathway.

### CheM signaling shifts SiiE from release to retention

The experiments described above excluded a motility-related effect to be responsible for elevated *Salmonella* invasion after MeAsp stimulation. Instead, the pronounced phenotype in conjunction with polarized cells and the identification of CheM as a SiiAB interaction partner argues for CheM as a regulator of SPI-4 dependent adhesion. Previous studies suggested that SPI-4 adhesion capability is determined by the amount and/ or binding strength of surface-localized SiiE (7, 14). Therefore, mechanisms regulating SiiE surface expression might account for SPI-4 dependent adhesion.

To test whether attractant binding to CheM affects localization of SiiE, we quantified the amounts of surface-retained and secreted SiiE adhesin after 3.5 h of growth (late logarithmic phase) with or without addition of MeAsp. Bacteria-associated SiiE was quantified in a dot blot assay using a SiiE-specific antibody and normalization to the LPS signal. Upon addition of MeAsp, elevated amounts of retained SiiE were detected for WT STM and the Δ7 MCP mutant carrying pCheM (Fig 5A). In contrast, no upregulation of surface-localized SiiE in response to MeAsp was observed for the Δ7 MCP mutant harboring the empty vector or for the Δ*siiE* mutant which served as negative control (Fig 5A). Quantification of secreted SiiE using a specific ELISA (6) revealed an inhibitory effect of CheM attractant binding reciprocal to SiiE surface localization. MeAsp inhibited SiiE secretion of WT and Δ7 MCP [pCheM] to the level of the Δ*siiE* control. Interestingly, compared to WT STM, almost 2-fold more SiiE was secreted from the Δ7 MCP strain harboring the empty vector control (Fig 5B).

**Fig 5.**
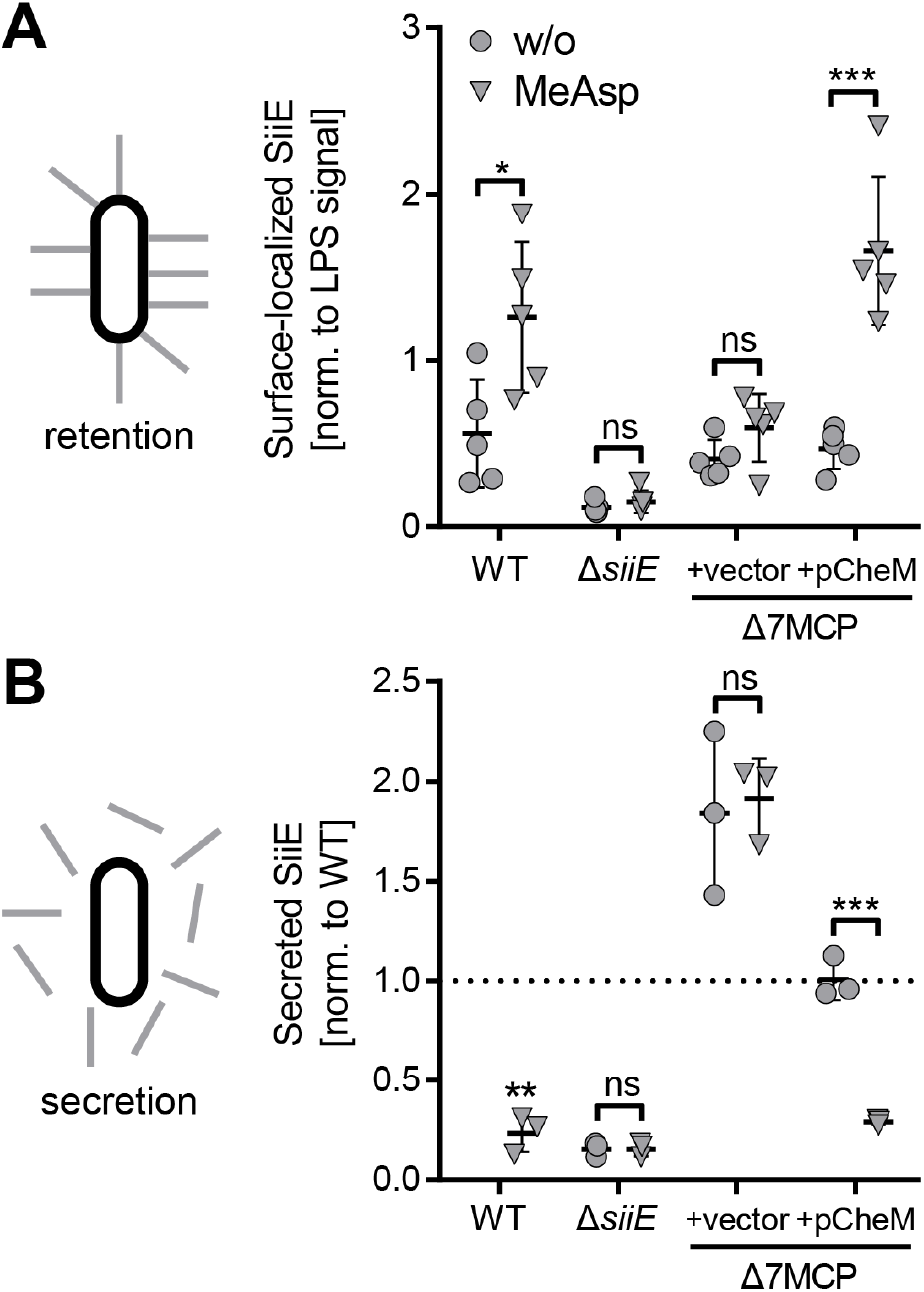
CheM-specific attractant binding promotes SiiE surface localization. (A) Dot-blots were used to quantify surface-localized SiiE and LPS (used for normalization) of *S*. Typhimurium (STM) wild-type (WT), a Δ*siiE* mutant or the Δ7 MCP deletion strain, transformed with the empty vector (vector) or pCheM. Bacteria were either grown without (w/o) or in the presence of 10 mM MeAsp. Mean data ± SD of five independent experiments are shown. (B) Secreted SiiE was quantified using an ELISA after 3.5 h of growth of the strains and under the conditions as described in (A). Mean data ± SD of three independent experiments done in triplicates are shown. Statistical significance was calculated using unpaired, two-tailed *t* test between groups or a one sample *t* test against the hypothetical value 1.0 (WT in (B)) and was defined as ns = not significant, * for *p* < 0.05, ** for *p* < 0.01 and *** for *p* < 0.001.

Our findings support a model where the interplay of CheM with the SPI-4 components SiiAB regulates SiiE localization. Attractant binding by CheM resulted in more surface-localized SiiE. To test whether indeed surface-retained SiiE can function as an adhesin, we combined competitive index experiments with a screen for ligand expression using immunomagnetic particles (SIMPLE) (37) (Fig 6A). The test and reference strains harboring different antibiotic resistance cassettes were mixed equally and an aliquot was plated on appropriate selective media to verify the proportion of the two strains. Subsequently, α-SiiE antibodies and magnetic protein A beads were added to the mixture. SiiE-positive bacteria were enriched through magnetic separation of beads coated with antibodies that have bound their antigen. Finally, the proportion of test and reference strain was determined through parallel plating (Fig 6A). Using STM WT as test strain and Δ*siiE* as reference, we achieved ∼8-fold enrichment using this assay. As expected, there was no effect of MeAsp on enrichment of STM WT over Δ*siiE* (Fig 6B). When the Δ7 MCP strain was used as reference, WT STM was ∼3-fold enriched in the presence of MeAsp, while no enrichment was seen without attractant (Fig 6B). These results demonstrate that addition of MeAsp enhanced the localization of SiiE on the surface of *Salmonella* in a MCP-dependent manner.

**Fig 6.**
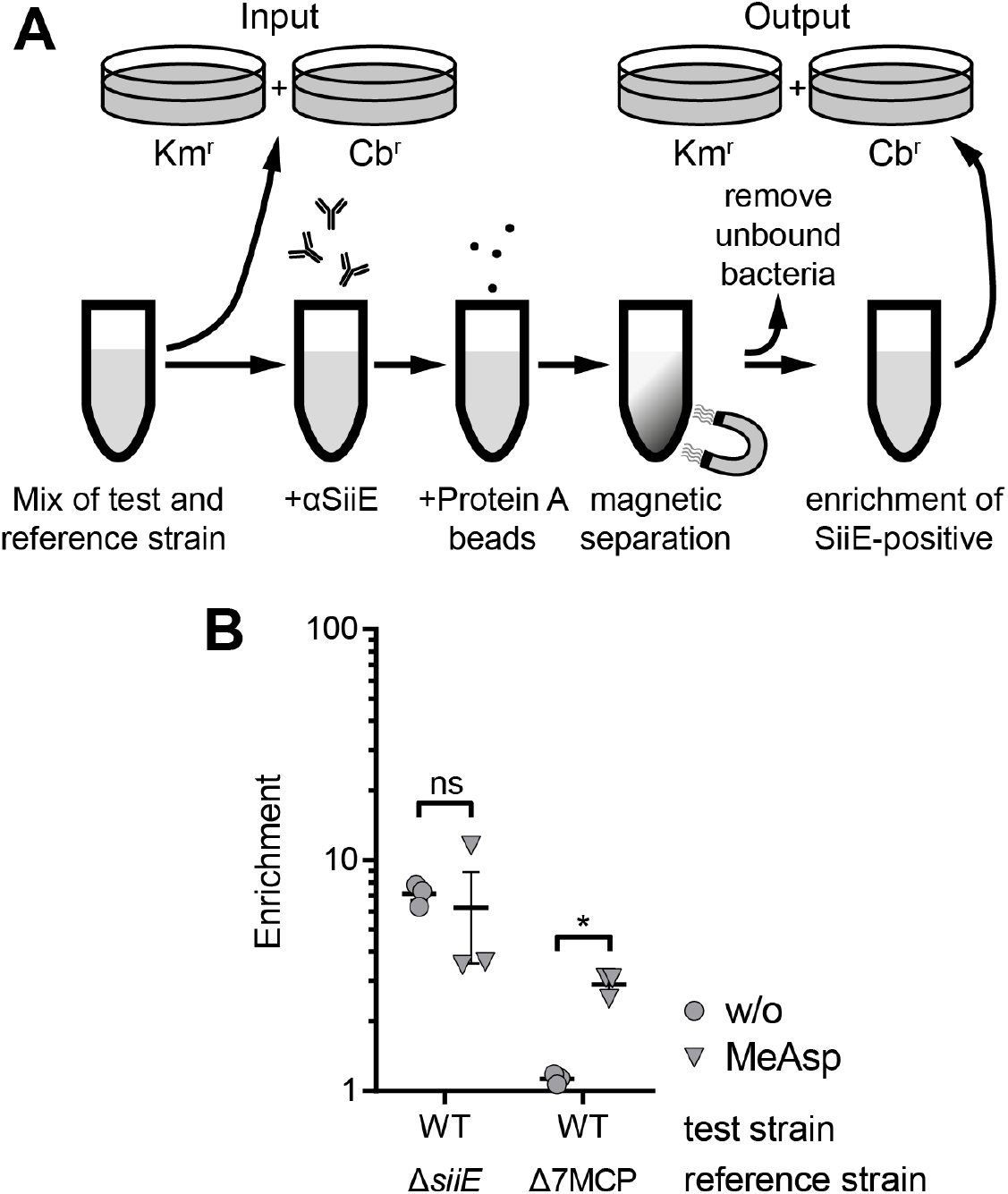
Increase of adhesion-competent SiiE in the presence of MeAsp. (A) Principle of a modified screening with immunomagnetic particles for ligand expression (SIMPLE) assay. (B) Enrichment of the test strains compared to the reference strains as indicated without (w/o) or in the presence of 10 mM MeAsp was determined using the SIMPLE assay as shown in (A). Data of three independent experiments done in triplicates are shown. Statistical significance was calculated using unpaired, two-tailed *t* test between groups as indicated and was defined as ns = not significant and * for *p* < 0.05.

## DISCUSSION

In the present study, we identified the MCP CheM as a novel SiiAB interaction partner and the binding of attractants to CheM as a positive regulator of SPI-4 dependent adhesion. We found that straight swimming *cheA* or *cheY* mutants, which resemble an attractant-bound “always off” state of the MCPs and are incapable of phosphoryl transfer, were attenuated for invasion in polarized cells. In contrast, straight swimming *Salmonella* showed a higher probability to invade other cell types (19, 25). Recently, the *Salmonella* MCP McpC was shown to promote a straight swimming phenotype that was dependent on the SPI-1 transcription factor HilD (38). When we bypassed the impact of motility and chemotaxis on bacteria-host cell interaction by centrifugation, addition of the CheM attractant MeAsp was still able to enhance invasion of polarized cells. This was particularly remarkable for the non-motile Δ*fliI* strain, which by itself rules out any involvement of the “classical” chemotaxis-motility pathway. In the absence of MeAsp, centrifugation of Δ*cheA* and Δ*cheY* mutants led to invasion rates comparable to WT. Strikingly, in the presence of the CheM attractant, these strains were hyperinvasive (∼20-25-fold of WT). In contrast, invasion capability was completely abolished with additional deletion of all 7 MCP genes. These observations cannot be explained with the chemotaxis phosphorelay signaling (18). Therefore, we propose a novel, non-canonical signal transduction from the MCP to SPI-4 encoded proteins resulting in increased adhesion and subsequent bacterial invasion. Links between chemotaxis and bacterial virulence functions are not unprecedented. In *Pseudomonas aeruginosa*, the putative MCP encoded by *PA2573* regulates genes involved in virulence and antibiotic resistance and the soluble chemoreceptor McpB was shown to be important for virulence in several infection models (39, 40). In *Cronobacter sakazakii*, a plasmid-encoded MCP was reported to have an impact on adhesion, invasion, motility and biofilm formation (41). The MCPs TcpI and AcfB of *Vibrio cholerae* were shown to be important for infant mouse colonization (42). For plant pathogenic bacteria such as *Agrobacterium tumefaciens* or *Xanthomonas oryzae* pv. *oryzae*, many chemoattractants can also act as virulence inducers (43, 44). However, in all these examples chemotaxis signaling is mechanistically linked to virulence through transfer of phosphoryl groups to alternative response regulators resulting in a virulence-specific transcriptional response (36).

The transduction of the CheM attractant signal is presumably based on the identified interaction of the MCP with the SPI-4 T1SS-associated SiiAB proton channel. It is tempting to postulate a regulation of SiiAB proton flux through direct interaction with attractant-bound CheM (Fig 7). The peptidoglycan (PG) binding domain of MotB was shown to function as a plug sealing the proton channel. Upon association with the flagellar motor complex, the MotB domain is shifted through PG binding and thereby enables proton flux and energy conversion of the system (45). Similarly, the SiiA PG binding domain (16) could be displaced from the SiiAB proton channel through interaction with attractant-bound CheM. In orthologous *E. coli* Tar dimers, Asp binding induces a piston-like movement of one alpha helix within the sensory domain. This movement is amplified in the cytosolic HAMP domains and finally transmitted to the hairpin tip bundle controlling CheA autokinase activity (18). In our model, the structural changes within the CheM ligand binding domain, and perhaps other portions of the molecule, would change the molecular interface between CheM and the SiiAB channel, thus enabling PG binding of SiiA and proton flux. The energy harvested from the transmembrane H^+^ gradient would then be transferred to the SPI-4 encoded T1SS by means of an energy-rich conformation resulting in retention of the SiiE molecule (Fig 7). Such energy transfer has been described for the SiiAB homologs ExbBD and TolQR inducing conformational changes in TonB and TolA, respectively (46, 47). Interestingly, *in vitro* studies with the isolated periplasmic domain of SiiA showed a pH-dependency of PG binding activity. SiiA PG-binding was observed at pH 5.8-6.2, but not between pH 6.7 and pH 8.0 (16). Here, slightly acidic pH as found in some parts of the gut could serve as another environmental signal to regulate the adhesion capacity of SPI-4. Alternatively, the observed pH-dependent PG binding could function as a proxy to ensure sufficient energization by detecting periplasmic protons contributing to the proton motif force (PMF). According to our model (Fig 7), *Salmonella* utilizes Asp as an environmental cue to control SiiE surface expression. Aspartate and other free amino acids are generated from oligopeptides originating from food through the activity of peptidases at the apical side of polarized enterocytes (48). The bulk of this amino acid liberation, and subsequent absorption, takes place in the proximal jejunum and is usually complete at the terminal ileum (48, 49). Although there is extensive catabolism of enteral Asp by enterocytes (50) and gut bacteria (51), the microbiota also releases free Asp through peptide degradation (52). Recently, Asp was found to contribute to initial murine gut colonization of STM by enabling hydrogen/fumarate-dependent anaerobic respiration. Aspartate is taken up in exchange of succinate by the high-affinity antiporter DcuABC and converted to the alternative electron acceptor fumarate (53). Therefore, the availability of Asp within the small intestine not only enables bacterial expansion in competition to the intestinal microbiota, but also contributes, amongst other environmental stimuli, to precise spatio-temporal control of bacterial adhesion to polarized epithelial cells.

**Figure 7.**
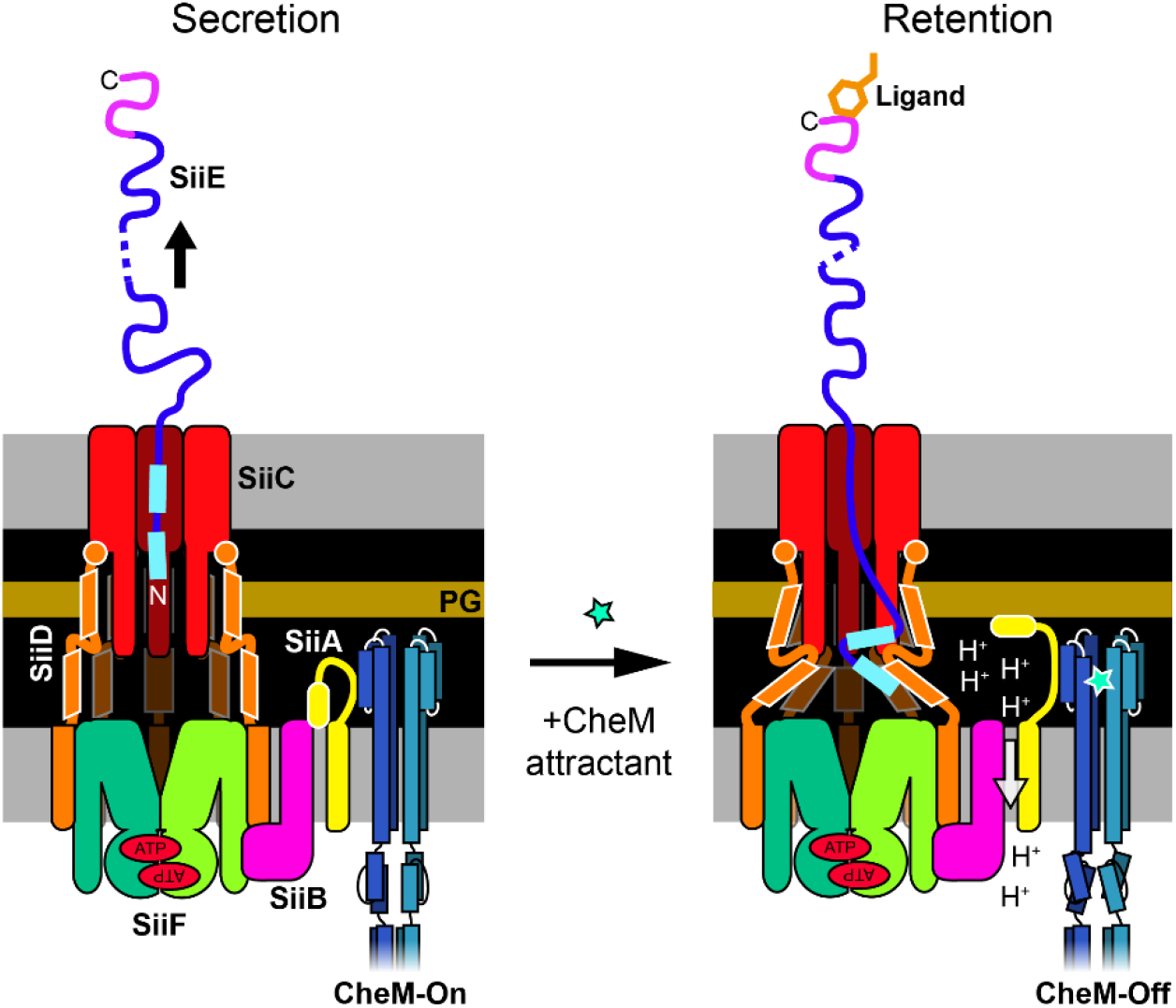
Proposed model how CheM could control SiiE-mediated adhesion. In the absence of attractant, CheM is in the kinase “on” state and the SiiAB proton channel is inactive, presumably due to the periplasmic peptidoglycan (PG) binding domain of SiiA functioning as a plug (left panel). Upon addition of CheM attractant, structural changes in the ligand binding domain of CheM induce displacement of the periplasmic SiiA portion and subsequent association with PG. Proton flux through SiiAB could energize structural changes in the T1SS thus retaining the N-terminal domain of SiiE within the channel (right panel).

In summary, we found that the MCP CheM interacted with the SPI-4 encoded SiiAB proton channel. Asp was identified as an attractant of CheM that elicited a change in the localization of the giant SPI-4-encoded adhesin SiiE of *Salmonella*: in the absence of Asp, SiiE was primarily secreted, whereas in the presence of Asp, SiiE was retained on the bacterial surface. Surface-bound, but not secreted SiiE functions as an adhesin. Thus, the CheM attractant L-aspartate acts as positive regulator of SPI-4-dependent adhesion to polarized cells. Although CheM directly interacts with the SPI-4 encoded SiiAB proton channel, the exact molecular mechanisms of signal transductions and adhesin retention remain to be characterized.

## MATERIALS AND METHODS

### Bacterial strains and plasmids

All strains used are listed in Table S1. Bacteria were routinely grown in LB media supplemented with 50 µg/mL carbenicillin (Cb) (Carl Roth, Mannheim, Germany), 25 µg/mL kanamycin (Km) (Carl Roth), 10 µg/mL chloramphenicol (Cm) (Carl Roth), 50 ng/mL anhydrotetracycline (AHT) (# 37919 Sigma-Aldrich, Schnelldorf, Germany), 10 mM α-aminoisobutyrate (AiBu) (#850993 Sigma-Aldrich) or 10 mM α-methyl-D, L-aspartate (MeAsp) (#M6001 Sigma-Aldrich), if required. For details on the construction of mutant strains and plasmids, the reader is referred to the supplementary information. Tables S2 and S3 give an overview of all the plasmids and primers used in this study, respectively.

### Protein-protein interaction assays

Bacterial two hybrid (B2H) assays were essentially carried out as described before (15). Briefly, *E. coli* reporter strain BTH101 was co-transformed with plasmids encoding for protein fusions with the T18 and T25 fragments of *Bordetella pertussis* CyaA. Transformants were spread on LB plates containing 25 µg/mL kanamycin, 100 µg/mL carbenicillin, 100 µM IPTG (Thermo Scientific, St. Leon-Rot, Germany) and 40 µg/mL X-Gal (5-bromo-4-chloro-3-indolyl-β-D-galactopyranoside; Thermo Scientific). Plates were incubated at room temperature for 48-72 h, and blue colonies indicated protein interaction resulting in functional CyaA-complementation.

For co-immunoprecipitation (co-IP), STM were either transformed with pWRG868 (CheM-3×Flag) or co-transformed with pWRG903 (CheM-HA) and pWRG905 (P_*tet*_::*siiAB*-3×Flag). O/N cultures were re-inoculated 1:31 into 500 mL fresh medium and grown with aeration for 3.5 h. The expression of SiiAB-3×Flag was induced for 2 h with AHT. Cells were pelleted (8,000 × g, 10 min, room temperature (RT)), re-suspended in 250 mL of pre-warmed MEM medium (Capricorn Scientific, Ebsdorfergrund, Germany) and allowed to grow for additional 30 min. Proteins were crosslinked with addition of 0.5% (w/v) paraformaldehyde (#43368 Alfa Aesar, Heysham, UK). After 15 min, the reaction was quenched with 0.125 M glycine. Cells were washed thrice with ice-cold MEM medium and then stored at −20 C. Pellets were re-suspended in 10 mL PBS supplemented with 1% *n*-dodecyl β-D-maltoside (# A0819, AppliChem, Darmstadt, Germany), 1× EDTA-free Halt protease inhibitor cocktail (#87785, Thermo Fisher Scientific, Karlsruhe, Germany), 0.5 mM MgSO_4_ and 5 µL TURBO DNase (Ambion). Subsequently, cells were lysed through a French pressure cell (EmulsiFlex-C3, Avestin, Mannheim, Germany) and debris was removed by low speed centrifugation (11,000 × g, 20 min, 4°C). The protein extract, containing either SiiB-3×Flag or CheM-3×Flag, was further cleared by additional centrifugation (20,000 × g, 15 min, 4°C). The protein concentration was measured by BCA assay (#23225, Thermo Fisher) and similar protein amounts were used for co-IP of all samples. Immunoprecipitation of 3×Flag-tagged proteins was performed using α-FLAG M2 affinity gel (Sigma-Aldrich) following the manufacturer’s recommendations. Therefore, 100 µL of the gel suspension (50 µL of packed gel volume) were washed 3× with 1 mL of PBS and subsequently added to each sample. Protein binding was allowed over night at 4°C. After three further washing steps, bound proteins were eluted from the beads with addition of 50 µL reducing sample buffer (Carl Roth) and heating for 15 min to 98°C.

### Western blot

Aliquots of protein samples were mixed with reducing sample buffer (Carl Roth) to a final concentration of 1×. After heating to 98°C for 15 min, 10 µL of each sample was analyzed by SDS-PAGE electrophoresis (ProSieve, Lonza, Cologne, Germany) and subsequent Western blot (Bio-Rad, Munich, Germany) on a polyvinylidene difluoride membrane (Thermo Fisher). Membranes were probed with antibodies against DnaK (clone 8E2/2, Enzo Life Science, Lörrach, Germany), SiiA, SiiB (15), HA (clone 3F10, Roche, Mannheim, Germany) or Flag (M2, Sigma-Aldrich) and appropriate horseradish-coupled secondary antibodies (Dianova, Hamburg, Germany).

### Mass spectrometry

STM was co-transformed with pWRG416 (P_*tet*_::*hilA*, resulting in mild SPI-1/4 overexpression) plus pWRG461 (*siiA*-3×Flag) or with pWRG416 plus pWRG462 (*siiAB*-3×Flag). Protein complexes were purified with α-FLAG M2 affinity gel (Sigma-Aldrich) as described for co-IP. After washing, bead-bound proteins were eluted twice with 450 µL of 0.1 M glycine pH3.5 and 180 µL of 0.5 M Tris-HCl pH7.4, 1.5 M NaCl was added for neutralization. To precipitate the proteins, trichloracetic acid was added to a final concentration of 10% and the samples were incubated O/N at 4°C. After centrifugation (20,000 × g, 45 min, 4°C), the pellets were washed twice with ice-cold acetone. The air-dried pellet was finally suspended in 100 µL fresh 50 mM NH_4_HCO_3_, pH7.8. The samples were subjected to tryptic digestion and the resulting peptide mixtures were analyzed by nano-ESI-LC-MS/MS (Thermo Scientific LTQ Orbitrap). Proteins were identified using Mascot (Matrix Science, London, UK) based on a custom proteome file of *S*. Typhimurium strain ATCC 14028s. Spectral counts were extracted using Scaffold Viewer (Proteome Software, Portland, OR, USA) and compared to controls (similar treated *S*. Typhimurium WT without 3×Flag tagged proteins) with the ‘R’ (54) package ‘apmsWAPP’ (55) applying upper quartile normalization and interquartile range filtering. Data is summarized in Table S4.

### Cell culture and infection

HT29-MTX human colonic epithelial cells (kind gift of G. Grassl, Hannover, Germany) were cultured in DMEM medium (high glucose, stable glutamine, sodium pyruvate) (Biowest, Nuaillé, France) supplemented with 10% FCS and non-essential amino acids (Biowest). HeLa cells (ATCC CCL-2, LGC Standards, Wesel, Germany) were grown in DMEM (Biowest) supplemented with 10% FCS, sodium pyruvate and 2 mM GlutaMax (Thermo Fisher) and MDCK cells (subclone Pf, Department of Nephroplogy, FAU Erlangen-Nürnberg) were kept in MEM medium (Biowest) supplemented with 10% FCS, 2 mM Glutamax (Thermo Fisher) and non-essential amino acids (Biowest). To each medium 100 U/mL penicillin and 100 μg/mL streptomycin (Biowest) were added. Cultures were incubated at 37°C in a humidified 5% (v/v) CO_2_ atmosphere. For invasion assays, HT29-MTX cells were seeded in 24-well culture plates (#662160, Cellstar, Greiner Bio-One, Frickenhausen, Germany) at a density of 4×10^4^ cells per well 21 days prior infection. MDCK and HeLa cells were seeded in 96-well plates (#655180, Greiner Bio-One) at a density of 8×10^4^ or 6×10^3^ per well, respectively. MDCK cells were allowed to polarize for 10–11 days. The culture medium was changed every other day and medium without antibiotics was used for the last medium change.

Gentamicin protection assays were essentially carried out as described previously [7]. Briefly, bacterial overnight (O/N) cultures grown in LB supplemented with appropriate antibiotics were re-inoculated 1:31 in fresh medium and grown aerobically for another 3.5 h. An inoculum corresponding to a multiplicity of infection (MOI) of 10 (HeLa) or 25 (MDCK, HT29-MTX) was prepared in MEM/DMEM and used to infect the cells for 25 min. After the cells had been washed thrice with PBS, MEM/DMEM containing 100 µg/mL gentamicin was applied to each well to kill remaining extracellular bacteria. After 1 h of incubation, the cell layers were washed again with PBS and then lysed for 10 min with PBS containing 1% Elugent (Merck Millipore, Darmstadt, Germany) and 0,0625% Antifoam B (Sigma-Aldrich, Schnelldorf, Germany) to liberate the intracellular bacteria. Serial dilutions of the inoculum and the lysates were plated on Mueller Hinton (MH) plates to determine the colony-forming units. Based on the inoculum the percentage of invasive bacteria was calculated and subsequently normalized to WT.

### Swarming assay

Swarming of different *Salmonella* strains was assessed on semi-solid agar LB plates (LB with 5 g/L NaCl, 0.5% agar) as described before (19). Briefly, a small amount (0.2 μL) of bacterial O/N cultures was applied onto the center of LB soft agar plate and incubated for six hours at 37°C. The diameters of the swarm colonies were measured and the plates were photographed.

### Capillary assay

Capillary assays were essentially performed as described before (28) with the following modifications: An U-shaped dam created from a piece of modelling clay and parafilm was mounted onto a microscopy slide. The chamber thus created was sealed with a cover slip and filled with 500 µL of a 3.5 h bacterial sub-culture. A 1 µL capillary (BLAUBRAND intraEND, Brand, Wertheim, Germany) was heat-sealed at one end and then filled with a 100 mM MeAsp solution. The capillary prepared in this way was immersed in the chamber for 1 h at 37°C. The capillary was then rinsed, the sealed end broken off and the capillary contents was emptied using a pipetting aid (Brand). Serial dilutions were plated and CFUs determined

### ELISA

Antisera were raised in rabbits against the recombinant C-terminal moiety of SiiE (7). For detection of SiiE, culture supernatants were filter-sterilized (0.45 µm syringe filters, Corning Life Sciences, Amsterdam, Netherlands), and aliquots of 50 µL were directly applied to 96-well Nunc MultiSorp microtiter plates (#467340 Thermo Fisher) overnight at 4°C in a humid chamber. The plates were washed three times with 200 µL/well of PBS supplemented with 0.05% Tween20 (PBS-Tween), and the rabbit anti-glutathione S-transferase(GST)-SiiE-C detection antibody diluted 1:1,000 in PBS plus 10% inactivated FCS (PBS-FCS) was applied for 2 h at RT. After five washes with PBS-Tween, 100 µL of the anti-rabbit horseradish peroxidase-coupled secondary antibody diluted 1:2,500 in PBS-FCS was added to each well for 30 min at RT. The wells were washed again seven times with PBS-Tween, and 50 µL of enzyme-linked immunosorbent assay (ELISA) horseradish peroxidase substrate (#555214, Becton Dickinson, Heidelberg, Germany) was added. After incubation in the dark at RT for 8 to 15 min, the reaction was stopped by the addition of 25 µL/well 0.5 M H_2_SO_4_ and A_450_ was measured using a M1000 plate reader (Tecan, Männedorf, Switzerland).

### Dot Blot

Bacterial strains were diluted 1:31 in LB from O/N cultures and grown at 37°C for 3.5 h. Aliquots of 1 mL of bacterial culture were collected, cells pelleted and re-suspended in 1 mL of sterile LB. After an additional washing step with sterile LB, bacterial suspensions were adjusted to OD_600_=1 in 500 μL of 3% PFA in PBS. After fixation of bacterial cells for 15 min at RT, cells were pelleted (10,000 × g, 5 min., RT) and re-suspended in 500 μL PBS. Five microliters of bacterial suspensions were spotted on nitrocellulose membrane pieces, set in a black 24-well plate, which have been pre-wetted with PBS and dried again before adding bacteria. After drying of the spots, membranes were blocked with 5% dry milk powder and 3% BSA in PBS/T (PBS + 0.1% Tween20) for at least 30 min. For detection of SiiE on the bacterial surface, antiserum against the C-terminal moiety of SiiE was diluted 1:5,000 in blocking solution and applied to the membrane. LPS was detected using antiserum against Salmonella O-antigen (Becton Dickinson) at the same dilution. After incubation O/N at 4°C, membranes were washed thrice with PBS/T and HRP-linked secondary antibody was added in a 1:50,000 dilution in PBS/T. After three additional washing steps with PBS/T, membranes were rinsed in PBS, substrate for HRP was applied and signals were quantified using a Tecan M1000 plate reader in luminescence mode.

### SIMPLE Assay

A screen for ligand expression using immunomagnetic particles (SIMPLE) assay was carried out as described by Nuccio *et al*. (37) with the following modifications. *Salmonella* strains were sub-cultured 1:31 from O/N for 3.5 h at 37°C and adjusted to OD_600_=2 in fresh PB buffer (=TN buffer (0.1 M Tris-HCl pH7.5, 0.15 M NaCl) plus 1% casein). Strains carried either plasmid pWSK29 or derivatives to exhibit carbenicillin resistance or plasmid pWSK129 for a kanamycin resistance. The strains were mixed at a ratio of 1:1 and bacteria were then pre-incubated with α-SiiE antibody or pre-immune serum in 650 µL PB-buffer for 1 h at RT with head-over-head rotation. Then, 100 µL of washed magnetic beads (BioMag Protein A, Qiagen, Hilden, Germany) resuspended in 100 µL TN-buffer were added to each sample (total volume of 750 µL) and incubated for two additional hours. After three washing steps with 750 µL TN buffer, beads were suspended in 1 mL PBS. Serial dilutions of all probes were plated in parallel on MH plates containing carbenicillin or kanamycin, CFU were determined and enrichment based on the competitive index and normalization to pre-immune serum controls was calculated.

## Acknowledgements

We thank Guntram Grassl (Hannover, Germany) for providing cell line HT29-MTX. This work was funded by grants of the German research foundation (www.DFG.de) to Y.A.M. (MU 1477 9/2), M.H. (HE 1964 13/2) and R.G.G. (GE 2533 2/2).

