## Supplemental Information for "A chemotactic sensor controls *Salmonella*-host cell interaction"

### Supplementary Information

### A chemotactic sensor controls *Salmonella* adhesion

#### Supplementary Materials and Methods

##### Strain construction

Mutants of *S*. Typhimurium NCTC 12023 (STM) were generated using an optimized λ Red recombination method with recombinase expression plasmid pWRG730 in combination with the kanamycin resistance cassette template plasmid pWRG717 as described before (1). In detail, primers SiiF626-scarless-for and SiiF628-scarless-rev were used to amplify the I-SceI *aph* cassette from pWRG717 and the resulting product was integrated in the STM wild-type (WT) genome resulting in strain WRG233. The cassette of WRG233 was replaced by the annealed oligonucleotides SiiF-E627Q-for and -rev generating strain WRG238 ($siiF^{E627Q}$). Similarly, *cheA* and *cheY* deletions were generated by amplifying the I-SceI *aph* cassette from pWRG717 using primers CheA-scarless-for and -rev or CheY-scarless-for and –rev, respectively. For *cheA*, the product was recombined in parallel in STM WT and strain WRG279 (Δ7 MCP) (1), resulting in strains WRG468 and WRG518. Primers CheA-cleandel-for and -rev were used to amplify a product from STM WT genomic DNA which was used in a second recombination step leading to strains WRG469 (Δ*cheA*) and WRG522 (Δ7 MCP Δ*cheA*). In case of *cheY*, the PCR product was recombined in WRG279 only resulting in strain WRG517. WRG517 was used for a second recombination step with a PCR product originating from primers CheY-cleandel-for and -rev with STM WT genomic DNA as template to generate WRG521 (Δ7 MCP Δ*cheY*). All mutant strains were verified by sequencing of the relevant genomic loci.

**Plasmid cloning**

Plasmid pWRG416 for tetracycline-inducible expression of HilA was cloned by amplifying the tetracycline promoter fragment from pST98-AS (2) with primers SmaI-XhoI-PtetA-for and PtetA-EcoRV-rev and the *hilA* gene from genomic DNA of STM WT using primers EcoRV-HilA-for and HilA-NotI-rev. After digestion of the fragments with EcoRV and EcoRV/NotI, respectively, both were cloned into PmlI/NotI-digested pETcoco-1 (Novagen, Darmstadt, Germany). Flag epitope-tagged variants of the *siiA* and *siiB* genes under control of the native *siiA* promoter ($P_{siiA}$) were cloned by amplifying $P_{siiA}$::*siiA* and $P_{siiA}$::*siiAB* from genomic STM WT DNA with primer Xho-SmaI-ProSPI4-for together with primer SiiA-NaeI-rev or SiiB-NaeI-rev. The 3×Flag tag was amplified from p3xFLAG-Myc-CMV-24 (Sigma-Aldrich, Schnelldorf, Germany) with primers EcoRV-3xFlag-for and 3xFlag-XbaI-rev. After digesting the *siiA* and *siiAB* fragments XhoI/NaeI and the 3×Flag fragment EcoRV/XbaI, each gene fragment and the epitope tag product was cloned in XhoI/XbaI-digested pWSK29 (3), resulting in pWRG461 and pWRG462. Plasmid pWRG674 was cloned by amplifying the *siiA* D13N mutated gene from plasmid pWRG648 (4) with primers PstI-p25-SiiA-for and SiiA-KpnI-rev which was subsequently digested PstI/Acc65I and ligated in similarly treated pKT25 (5). To construct pWRG676, *siiD* was PCR-amplified from STM WT genomic DNA using primers XbaI-p25-SiiD-for and SiiD-KpnI-rev and the fragment was cloned in pKT25 via KpnI/XbaI. In plasmids pWRG844 and pWRG845 *cheM* is fused to the *t25* or *t18* fragment, respectively. For construction, the *cheM* gene was amplified from the STM WT genome using primers XbaI-CheM-for and CheM-KpnI-rev and subsequently cloned via XbaI/KpnI in p25-N (6) and pUT18 (5). Plasmid pWRG846 was constructed by cloning the *cheM* fragment in pKT25 (5) resulting in a fusion of *t25* to *cheM*.

For construction of all remaining plasmids assembly cloning of PCR fragments was used (7). For C-terminal 3×Flag epitope-tagging of CheM, the *cheM* gene including its promoter

($P_{cheM}$) was amplified from the STM WT chromosome with primers pWSK-CheM-Gbs-for and CheM-3xFlag-Gbs-rev. Primers CheM-3xFlag-Gbs-for and pWSK29-Gbs-rev were used to linearize pWSK29 and assembly cloning resulted in pWRG868. In plasmids pWRG889 and pWRG890 the STM *cheM* gene present in pWRG847 (1) and pWRG868 was exchanged by *E. coli tar*. For pWRG889 the *tar* gene was amplified from *E. coli* K-12 DNA using primers PcheM-Tar-Gbs-for and Tar-pWSK-Gbs-rev and plasmid pWRG847 was used as PCR template with primers pWSK29-Gbs-for and PcheM-Gbs-rev. For pWRG890 the *tar* gene was amplified from *E. coli* K-12 DNA using primers PcheM-Tar-Gbs-for and Tar-3xFlag-Gbs-rev and plasmid pWRG868 was used as PCR template with primers 3xFlag-Gbs-for and PcheM-Gbs-rev. For HA epitope-tagging of CheM, the gene was amplified from STM WT DNA using primers pWSK-CheM-Gbs-for and CheM-HA-Gbs-rev. Primers CheM-HA-Gbs-for and pWSK29-Gbs-rev were used to linearize pWSK29 and assembly cloning resulted in pWRG903. Plasmid pWRG905 allows for tetracycline-inducible expression of SiiAB-3×Flag and was cloned as follows: The backbone plasmid pET-coco1 including a tetracycline-inducible promoter was amplified from pWRG416 with primers pETcoco-Gbs-for and RBS-PtetA-Gbs-rev. The $P_{siiA}$::*siiAB*::3×Flag fragment was amplified from pWRG462 using primers PtetA-SiiA-Gbs-for and 3xFlag-pETcoco1-Gbs-rev which was finally assembled with the backbone fragment to yield pWRG905. All plasmids were verified by analytical restriction digests and subsequent sequencing of the cloned regions.

**Staining and microscopy**

HT29-MTX cells were seeded in 24-well culture plates (#662160, Cellstar, Greiner Bio-One, Frickenhausen, Germany) at a density of $4\times10^4$ cells per well one or 21 days prior Periodic acid-Schiff (PAS) staining. The cells were maintained in DMEM medium (high glucose, stable glutamine, sodium pyruvate) (Biowest, Nuaillé, France) supplemented with 10% FCS and non-essential amino acids (Biowest). Mucins were visualized using a PAS staining kit

(#101646 Sigma-Aldrich, Schnelldorf, Germany) according to manufacturer's instructions but omitting the hematoxylin staining, dehydration and xylene clearing. Stained wells were imaged using an Eclipse Ti-E microscope (Nikon) with 20× objective. Image capturing was done with NIS Elements Ar software (v4.2) from a DS-Fi1 color camera (Nikon).

#### Supplementary Figures

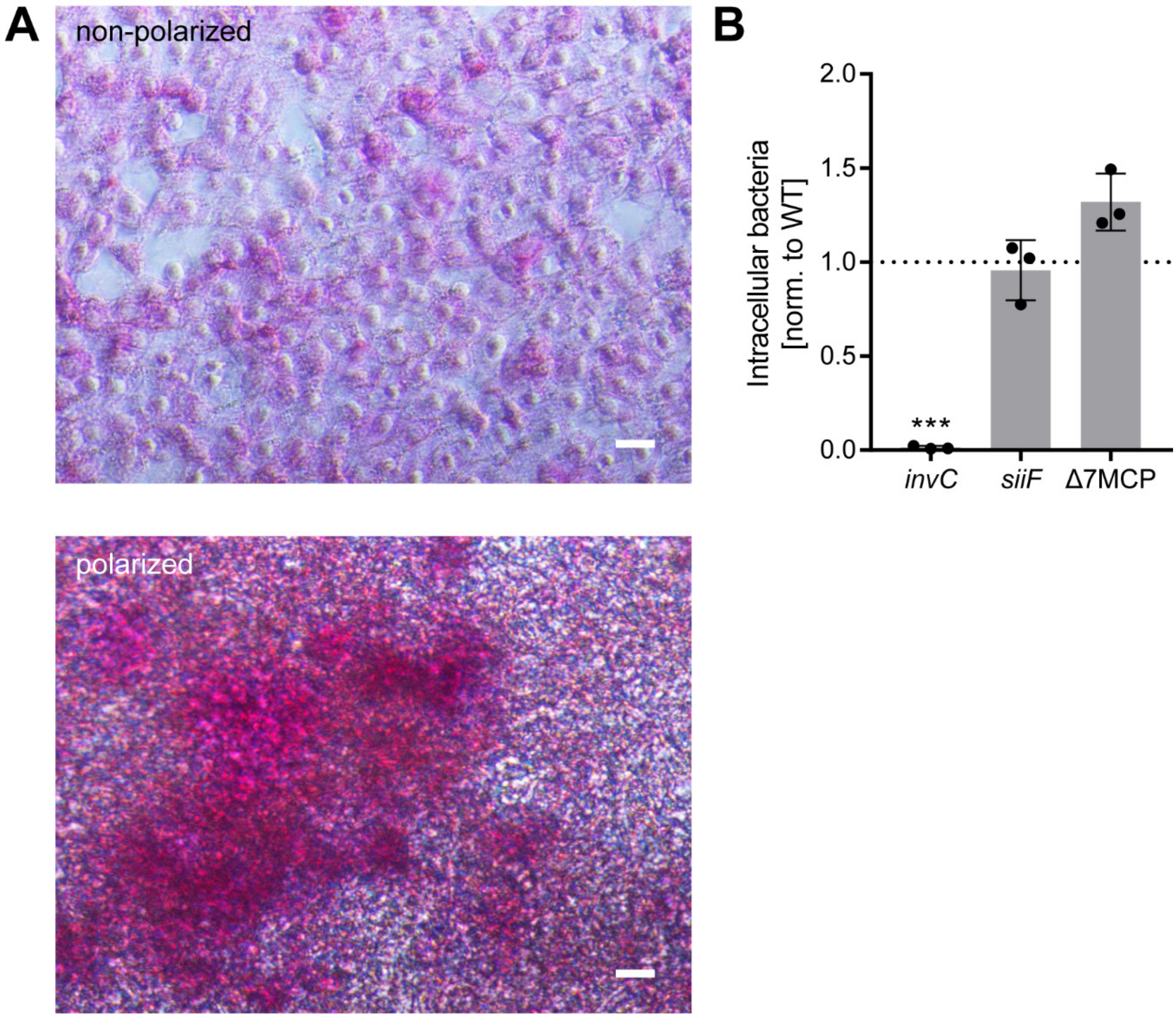


**Figure S1. Characterization and infection of HT29-MTX cells.** (A) Periodic acid-Schiff (PAS) staining of mucins (purple) produced by HT29-MTX after one day (non-polarized, upper panel) and after 21 days (polarized, lower panel) of differentiation. Scale bar = 20 µm. (B) Relative invasion rates in non-polarized HT29-MTX cells as normalized to *S.* Typhimurium (STM) wild-type (WT) (black dotted line) after one hour of infection with strains *invC* (non-functional T3SS-1), *siiF* (non-functional SPI-4) and Δ7 MCP are depicted. Statistical significance was calculated using a one sample *t* test against the hypothetical value 1 and were defined as *** for $p < 0.001$.

#### Supplementary Tables

**Table S1. Strains used in this study.**

| Strain | Relevant characteristic(s) | Source or Reference |
|---|---|---|
| *E. coli* strains | | |
| BTH101 | F-, cya-99, araD139, galE15, galK16, rpsL1 (Str$^r$), hsdR2, mcrA1, mcrB1 | (8) |
| *Salmonella* strains | | |
| MvP589 | NCTC 12023 ΔSPI-4 FRT | (9) |
| MvP593 | NCTC 12023 Δ*siiB* FRT | (9) |
| MvP599 | NCTC 12023 Δ*siiE* FRT | (9) |
| MvP771 | NCTC 12023 Δ*siiA* FRT | (10) |
| MvP818 | NCTC 12023 Δ*invC* FRT | (11) |
| MvP1212 | NCTC 12023 Δ*cheY* FRT | (12) |
| MvP1213 | NCTC 12023 Δ*fliI* FRT | (13) |
| NCTC 12023 | wild type (**WT**), Nal$^s$, isogenic to ATCC 14028 | NCTC, Colindale, UK |
| WRG233 | NCTC 12023 *siiF626*::I-SceI *aph*, Km$^r$ | This study |
| WRG238 | NCTC 12023 *siiF*$^{E627Q}$, non-functional Walker B motif | This study |
| WRG246 | NCTC 12023 Δ*aer* | (1) |
| WRG255 | NCTC 12023 Δ*aer* Δ*tcp* | (1) |
| WRG260 | NCTC 12023 Δ*aer* Δ*tcp* Δ*tsr* | (1) |
| WRG264 | NCTC 12023 Δ*aer* Δ*tcp* Δ*tsr* Δ*trg* | (1) |
| WRG269 | NCTC 12023 Δ*aer* Δ*tcp* Δ*tsr* Δ*trg* Δ*cheM* | (1) |
| WRG277 | NCTC 12023 Δ*aer* Δ*tcp* Δ*tsr* Δ*trg* Δ*cheM* Δ*mcpC* | (1) |
| WRG279 | **Δ7 MCP**; NCTC 12023 Δ*aer* Δ*tcp* Δ*tsr* Δ*trg* Δ*cheM* Δ*mcpC* Δ*mcpB* | (1) |
| WRG468 | NCTC 12023 *cheA*::I-SceI *aph*, Km$^r$ | This study |
| WRG469 | NCTC 12023 Δ*cheA* | This study |
| WRG517 | NCTC 12023 Δ*aer* Δ*tcp* Δ*tsr* Δ*trg* Δ*cheM* Δ*mcpC* Δ*mcpB* *cheY*::I-SceI *aph*, Km$^r$ | This study |
| WRG518 | NCTC 12023 Δ*aer* Δ*tcp* Δ*tsr* Δ*trg* Δ*cheM* Δ*mcpC* Δ*mcpB* *cheA*::I-SceI *aph*, Km$^r$ | This study |
| WRG521 | **Δ7 MCP Δ*cheY***; NCTC 12023 Δ*aer* Δ*tcp* Δ*tsr* Δ*trg* Δ*cheM* Δ*mcpC* Δ*mcpB* Δ*cheY* | This study |
| WRG522 | **Δ7 MCP Δ*cheA***; NCTC 12023 Δ*aer* Δ*tcp* Δ*tsr* Δ*trg* Δ*cheM* Δ*mcpC* Δ*mcpB* Δ*cheA* | This study |

**Table S2. Plasmids used in this study.**

| Plasmid | Relevant characteristic(s) | Source or Reference |
|---|---|---|
| p25-N | encodes the T25 fragment (aa 1–224) of CyaA for C-terminal fusions; Km$^r$ | (6) |
| pKD4 | *aph* resistance cassette flanked by FRT sites, λ Pir dependent replication, Km$^r$, Ap$^r$ | (14) |
| pKNT25-ZIP | pKT25 derivative; leucine zipper of GCN4 fused in frame to T25 fragment; Km$^r$ | (5) |
| pKT25 | encodes the T25 fragment (aa 1–224) of CyaA; for N-terminal fusions; Km$^r$ | (5) |
| pUT18 | T18 fragment (aa 225–399) of CyaA; for C-terminal fusions, Ap$^r$ | (5) |
| pUT18C-ZIP | pUT18C derivative; leucine zipper of GCN4 fused in frame T18 fragment, Ap$^r$ | (5) |
| pWRG717 | pBluescript II SK+ derivative, *aph* resistance cassette and I-SceI cleavage site, Km$^r$, Ap$^r$ | (1) |
| pWRG730 | pSIM5 (15) derivative, temperature-sensitive replication (30°C) and Red recombinase expression (42°C), Tet-inducible expression of I-SceI, Cm$^r$ | (1) |
| pWRG416 | P$_{tet}$::*hilA* in pETcoco-1, Cm$^r$ | This study |
| pWRG461 | P$_{siiA}$::*siiA*-3×Flag, Ap$^r$ | This study |
| pWRG462 | P$_{siiA}$::*siiAB*-3×Flag, Ap$^r$ | This study |
| pWRG551 | P$_{tac}$::*siiF*-T25 in p25N, Km$^r$ | (4) |
| pWRG554 | P$_{tac}$::*siiB*-T25 in p25N, Km$^r$ | (4) |
| pWRG643 | P$_{tac}$::T25-*siiA* in pKT25, Km$^r$ | (4) |
| pWRG674 | P$_{tac}$::T25-*siiA*-D13N in pKT25, Km$^r$ | This study |
| pWRG676 | P$_{tac}$::T25-*siiD* in pKT25, Km$^r$ | This study |
| pWRG844 | P$_{tac}$::*cheM*-T25 in p25N, Km$^r$ | This study |
| pWRG845 | P$_{tac}$::*cheM*-T18 in pUT18, Ap$^r$ | This study |
| pWRG846 | P$_{tac}$::T25-*cheM* in pKT25, Km$^r$ | This study |
| pWRG847 | **pCheM**; P$_{cheM}$::*cheM* in pWSK29, Ap$^r$ | (1) |
| pWRG868 | P$_{cheM}$::*cheM*-3×Flag in pWSK29, Ap$^r$ | This study |
| pWRG889 | **pTar**; P$_{cheM}$::*tar* in pWSK29, Ap$^r$ | This study |
| pWRG890 | P$_{cheM}$::*tar*-3×Flag in pWSK29, Ap$^r$ | This study |
| pWRG903 | P$_{cheM}$::*cheM*-HA in pWSK29, Ap$^r$ | This study |
| pWRG905 | P$_{tet}$::*siiAB*-3×Flag in pETcoco-1, Cm$^r$ | This study |
| pWSK29 | Low-copy-number vector, Ap$^r$ | (3) |
| pWSK129 | Low-copy-number vector, Km$^r$ | (3) |

**Table S3. Primers used in this study.**

| Primer name | Sequence 5'→ 3' |
|---|---|
| 3xFlag-Gbs-for | GACTACAAAGACCATGACGG |
| 3xFlag-pETcoco-Gbs-rev | TCTTCCGGAGCGAGTTCTGGCTGGCTTGCGTTACTTGTCATCGTCATCCTTGTAATC |
| 3xFlag-XbaI-rev | GCCTCTAGATTACTTGTCATCGTCATCCTTGT |
| CheA-cleandel-for | ACATTACTCATACCGGTCATATTATTCCCTTCTCACTCAAGCTATCACCTCGGTTCCGCT |
| CheA-cleandel-rev | ACCGATGACTTTCCCTAC |
| CheA-scarless-for | ACATTACTCATACCGGTCATATTATTCCCTTCTCACTCAAAGGGTTTTCCCAGTCACGAC |
| CheA-scarless-rev | AGCGTACCCACATCGCCAAAAGCGGAACCGAGGTGATAGCTGCTTCCGGCTCGTATGTTG |
| CheM-3xFlag-Gbs-for | GACTACAAAGACCATGACGGTGATTATAAAGATCATGACATCGATTACAAGGATGACGATGACAAGTGATCGACGTGCGCTGTCGG |
| CheM-3xFlag-Gbs-rev | CTTGTCATCGTCATCCTTGTAATCGATGTCATGATCTTTATAATCACCGTCATGGTCTTTGTAGTCGAAGGTTTCCCAGTTCGCAT |
| CheM-HA-Gbs-for | TACCCATACGACGTCCCAGACTACGCTTGATCGACGTGCGCTGTCGG |
| CheM-HA-Gbs-rev | AGCGTAGTCTGGGACGTCGTATGGGTAGAAGGTTTCCCAGTTCGCATC |
| CheM-KpnI-rev | CTTAGGTACCAAGGTTTCCCAGTTCGCATCATCCCTGGCGGC |
| CheY-cleandel-for | CCGGACAGGCGATACGTATTTGAACCAGGAGTAGTATTTTGGATGCGATGATGATGCAAC |
| CheY-cleandel-rev | ACGCCAATAGGCAGAGTAAG |
| CheY-scarless-for | CCGGACAGGCGATACGTATTTGAACCAGGAGTAGTATTTTAGGGTTTTCCCAGTCACGAC |
| CheY-scarless-rev | ATCAGCAGGCTTGATAGATGGTTGCATCATCATCGCATCCTGCTTCCGGCTCGTATGTTG |
| EcoRV-3xFlag-for | CAGGATATCGACTACAAAGACCATGACGG |
| EcoRV-HilA-for | GGCGATATCATGCCACATTTTAATCCTG |
| HilA-NotI-rev | TGCGCGGCCGCTTACCGTAATTTAATCAAGCGGG |
| PcheM-Gbs-rev | AAGGCACCTTCCTGATAACG |
| PcheM-Tar-Gbs-for | TGCCGATAACGTTGATAACTCGTTATCAGGAAGGTGCCTTATGATTAACCGTATCCGCGT |
| pETcoco-Gbs-for | CGCAAGCCAGCCAGAACTCGCTCC |
| PstI-p25-SiiA-for | GACCTGCAGTGGAAGACGAAAGTAATCCG |
| PtetA-EcoRV-rev | GCGGATATCTTTCTCTATCACTGATAGGGAG |
| PtetA-SiiA-Gbs-for | CCACTCCCTATCAGTGATAGAGAAAGATATCATGGAAGACGAAAGTAATCCG |
| pWSK29-Gbs-for | TAATTGCGCGCTTGGCGTAATC |
| pWSK29-Gbs-rev | ACCCAATTCGCCCTATAGTG |
| pWSK-CheM-Gbs-for | TAATACGACTCACTATAGGGCGAATTGGGTCGTCGTCGGGATAGTGGTAG |
| RBS-PtetA-Gbs-rev | GATATCTTTCTCTATCACTGATAGG |
| SiiA-KpnI-rev | CTTAGGTACCTCTGACACCTTTTTATTAATAG |
| SiiA-NaeI-rev | TACGCCGGCCTCTGACACCTTTTTATTAA |
| SiiB-NaeI-rev | TACGCCGGCATCTTCATTTTTTTCCTCCTTG |
| SiiD-KpnI-rev | CTTAGGTACCCAAGGTGTATCTAATCGTTTAG |
| SiiF626-scarless-for | TATTATTAGCACGTAGTCTGAGTAGTGACGCCAGCGTCTTTTTATGGGATAGGGTTTTCCCAGTCACGAC |
| SiiF628-scarless-rev | AAGTTATCAAAAATTTGCTTCTCGGTATTCTCATCCAGATTTGATGTTGGTGCTTCCGGCTCGTATGTTG |
| SiiF-E627Q-for | CGTAGTCTGAGTAGTGACGCTAGCGTCTTTTTATGGGATCAGCCAACATCAAATCTGGATGAGAATACCGAGAAGCAAAT |
| SiiF-E627Q-rev | ATTTGCTTCTCGGTATTCTCATCCAGATTTGATGTTGGCTGATCCCATAAAAAGACGCTAGCGTCACTACTCAGACTACG |

| | |
|---|---|
| SmaI-XhoI-PtetA-for | ATTCTCGAGCCCGGGTCGATGGGTGGTTAACTC |
| Tar-3xFlag-Gbs-rev | TGTCATGATCTTTATAATCACCGTCATGGTCTTTGTAGTCAAATGTTTCCCAGTTTGGAT |
| Tar-pWSK-Gbs-rev | ATTCGCCCTTATGACCATGATTACGCCAAGCGCGCAATTATCAAAATGTTTCCCAGTTTG |
| XbaI-CheM-for | CCGTCTAGAGTTTAACCGTATCCGCGTTGTCACAATGC |
| XbaI-p25-SiiD-for | GACTCTAGAGAATAGAAGACAAAGCGATCATC |
| XhoI-SmaI-ProSPI4-for | GATCTCGAGCCCGGGAAAGCGTTATTTGCATTTTCG |
